# Interferometric scattering microscopy reveals microsecond nanoscopic protein motion on a live cell membrane

**DOI:** 10.1101/401133

**Authors:** Richard W. Taylor, Reza Gholami Mahmoodabadi, Verena Rauschenberger, Andreas Giessl, Alexandra Schambony, Vahid Sandoghdar

## Abstract

Much of the biological functions of a cell are dictated by the intricate motion of proteins within its membrane over a spatial range of nanometers to tens of micrometers and time intervals of microseconds to minutes. While this rich parameter space is not accessible to fluorescence microscopy, it can be within reach of interferometric scattering (iSCAT) particle tracking. Being sensitive even to single unlabeled proteins, however, iSCAT is easily accompanied by a large speckle-like background, which poses a substantial challenge for its application to cellular imaging. Here, we show that these difficulties can be overcome and demonstrate tracking of transmembrane epidermal growth factor receptors (EGFR) with nanometer precision in all *three dimensions* at up to microsecond speeds and tens of minutes duration. We provide unprecedented examples of nanoscale motion and confinement in ubiquitous processes such as diffusion in the plasma membrane, transport on filopodia, and endocytosis.

## Introduction

Recent steady and rapid progress in fluorescence microscopy has brought us closer to monitoring cellular events with nanometer spatial resolution, however, there still remains a great deal to accomplish ^1^. A major challenge is that the finite emission rate of a fluorescent source such as a dye molecule or a semiconductor quantum dot precludes high localization precision within microseconds, simply because too few photons are emitted in very small windows of time. Fluorescent probes also succumb to photo-degradation which heavily restricts the duration of observation. To overcome these limitations, Rayleigh scattering from noble metal nanoparticles has been detected via dark-field, differential interference contrast or bright-field microscopy ^2–5^. A central difficulty in scattering-based microscopy is that the contribution from a nanoscopic probe competes against background scattering and a low signal-to-noise ratio (SNR). As a result, such efforts have been limited in positional uncertainty to several tens of nanometers in high-speed tracking experiments ^6^, unless large probes were used ^5^.

In this work, we employ interferometric scattering (iSCAT) microscopy ^7–9^ to track proteins in live cell membranes, presenting a substantial spatial and temporal leap over the impressive fluorescence-based developments in visualizing the interaction of a probe with the cell and the association between diffusional dynamics and local topology ^10^. Here, we use gold nanoparticles (GNP) to label epidermal growth factor receptors (EFGR) in HeLa cells. EGFR is a single-pass type I transmembrane protein and a member of the signaling receptor tyrosine kinase family, which senses and responds to extracellular signals and whose aberrant signaling has been linked to a variety of cancers^11,12^. In what follows, we provide examples of sub-diffusion and nanoscopic confinement in the *three-dimensional* (3D) motion of a protein at very high temporal resolution and for long durations. Furthermore, we show that a GNP-labeled protein can be used as a *nano-rover* to map the nanoscopic topology of cellular features such as membrane terrains, filopodia or clathrin structures.

### Three-dimensional iSCAT microscopy

The principle of iSCAT microscopy is that light scattered from a nanoparticle of interest is interfered with a brighter reference field, usually one reflected from the sample substrate ^7^. The high SNR and sensitivity of iSCAT has allowed fast detection and tracking of very small nano-objects down to single unlabeled proteins, but this has been achieved in model synthetic systems with minimal background scattering such as clean substrates ^13–15^, supported bilayers ^16–18^ or giant unilamellar vesicles ^19^. When iSCAT microscopy is applied to the membrane of the cell, the local roughness of the plasma membrane and the nanoscopic heterogeneity of the cellular corpus give rise to a dynamic speckle-like background, which hinders detection and tracking of small nano-objects.

Details about iSCAT microscopy, probe functionalization and cell culturing are provided in **Methods**. Here, it suffices to mention that each GNP (diameter 48 nm) was functionalized with epidermal growth factor (EGF) proteins and we verified the probes stimulate the EGFR. This size of probe is reported to introduce negligible effects from interactions with the crowded cellular environment ^20^. As illustrated in **Fig. 1a**, EGF-GNP probes were introduced to the sample chamber on the microscope via a micropipette to label endogenous EGFR on HeLa cells. **Figure 1b** shows an iSCAT image of a part of a HeLa cell upper surface before GNPs were added. The membrane surface appears to be corrugated with speckle-like contrast variations up to 50% and spatial features as small as the diffraction limit. The challenge of live-cell iSCAT microscopy is to account for this dynamic background and extract the point-spread function (PSF) of the nanoparticle under study in order to reconstruct its 3D trajectory.

**Figure 1:**
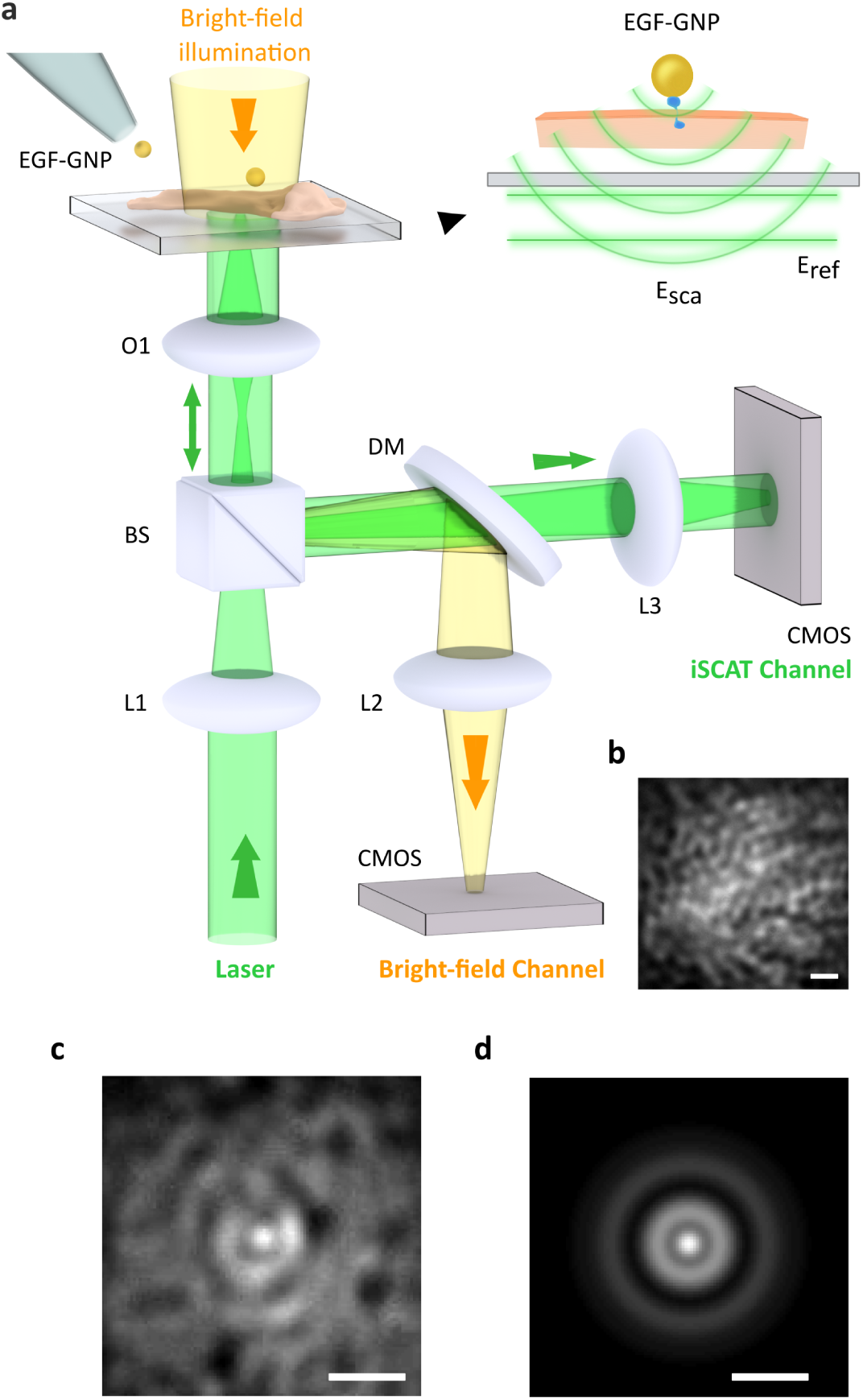
(a) Experimental arrangement of the iSCAT microscope for live-cell imaging. Cells are plated in a glass-bottomed dish under Leibowitz medium. A micropipette delivers the EGF-GNP probes directly onto the cell culture, where they specifically target the EGFR protein in the cell membrane. The bright-field illumination channel from above assists in inspecting the culture but is not required for iSCAT imaging. L1-L3: lenses, O1: 100x objective, BS = 90:10 beam splitter, DM = 590 nm short-pass dichroic mirror. iSCAT imaging was performed with illumination intensities of 1-8 kWcm^*-*2^, which is known to be viable for HeLa at our wavelength ^24^. Inset: wavefronts of the fields contributing to the iSCAT signal. (b) A section of the membrane of the HeLa cell before labeling viewed via reflection iSCAT. (c) iSCAT image of the cell membrane including a bound EGF-GNP probe. (d) PSF extracted from (c). Scale bars in (b-d) are 1 *µ*m.

In **Fig. 1c**, we present an example of a snapshot of a GNP bound to the plasma membrane, yielding a positive iSCAT contrast. To determine the position of the GNP with nanometer precision, we have developed an algorithm to extract the iSCAT-PSF, which is composed of a series of concentric rings caused by the interference of plane and spherical waves on the camera (see **Fig. 1a**). **Figure 1d** displays the outcome for the example presented in **Fig. 1c**. The interferometric nature of the iSCAT signal also leads to contrast modulations as the scatterer changes its axial position. This feature has been used to track small axial movements by monitoring the brightness of the central spot of the PSF^19,21^. However, the periodicity of this contrast introduces directional ambiguity in longer trajectories. In this work, we address this issue by analyzing the separations and amplitudes of the PSF rings, which posses height-specific signatures ^22^. The large SNR in our measurements results in an exquisite localization precision of *σ* = 2-6 nm for both the lateral and axial directions, taking us beyond conventional 2D trajectories that can be commonly found in the literature ^23^.

## Results

The spatio-temporal dynamics of protein function in the membrane as well as its uptake and trafficking are expected to be strongly influenced by heterogeneous interactions with various cellular components such as actin filaments, clathrin structures, other membrane proteins, and the extracellular matrix ^25^. A lack of means to directly visualize these features at high spatial and temporal resolution has left much of their details a matter of debate. We now discuss numerous trajectories, which reveal a wealth of dynamic 3D heterogeneities and shed new light on protein motion.

### Quantitative study of sub-diffusion in the plasma membrane

We begin by considering a 2D example of the EGFR journey on the plasma membrane of a living HeLa cell. **Figure 2a** presents a trajectory that results from a video containing *N*=75,000 frames over 3.75 s recorded at 20,000 frame per second (fps). The high SNR yields *σ* = 6 nm lateral spatial precision within the frame exposure time of 17.5 *µ*s. A visual inspection of the trajectory, which is temporally color coded, indicates a strong heterogeneity.

**Figure 2:**
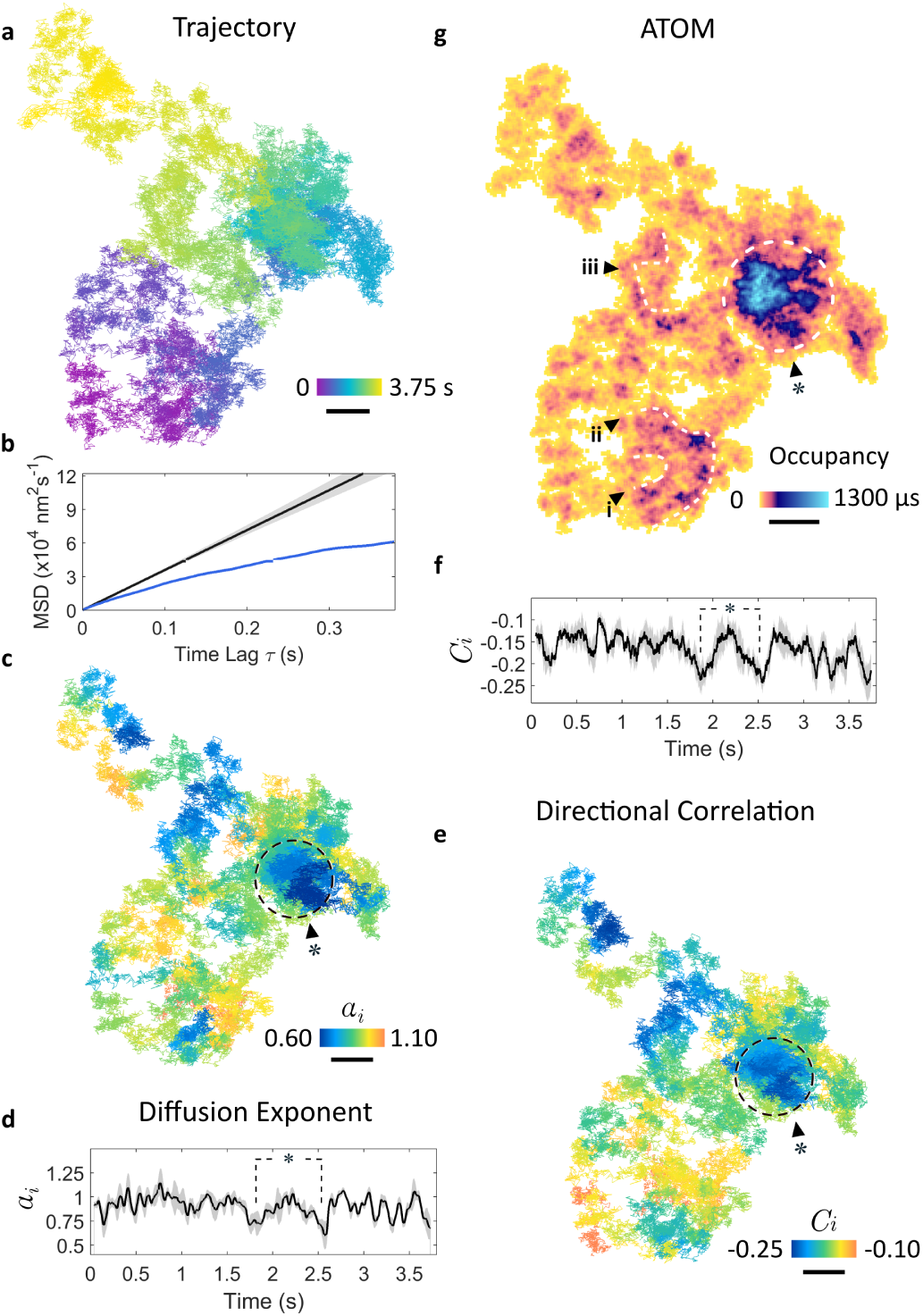
Diffusion on the plasma membrane, resolved with 17.5 *µ*s temporal resolution. (a) Lateral diffusional trajectory where time is encoded by color. (b) The blue curve shows the global mean square displacement of the trajectory in (a). The black linear curve is a simulation of normal diffusion (*α* = 1). Shadowed region indicates the uncertainly in the simulated MSD. (c) Diffusional exponent parameter *α*_*i*_ of the *i*-th rolling window plotted over the lateral trajectory. Regions of sub-diffusion in which *α <* 1 are pronounced through the choice of the color code. (d) *α*_*i*_ as a function of time. The gray shadowed region presents a mean uncertainty of 7±4%, corresponding to a 95% confidence interval for the fitted *α*_*i*_ extracted for a window of 100 ms (1000 frames) and time lag intervals of *τ* = 250 *µ*s. Points marked with * correspond to the region of dashed circle in (c). (e) Step-direction correlation function *C*_*i*_ for small rolling windows along the trajectory. (f) Step-direction correlation plotted as a function of time. Shadowed regions denote the uncertainty in *C*_*i*_. (g) ATOM occupation plot. The areal bin size was set to that of the localization error. Noteworthy regions of extended occupation are marked with annotations. The enclosed region represents a spatially dense patch of notable sub-diffusion. Scale bar denotes 100 nm throughout.

To analyze the diffusion behavior in a quantitative fashion, we first compute the mean square displacement (*MSD*) for the whole trajectory, employing the conventional formula of *MSD* = 4*Dτ* + 2*σ*^2^, where *D* is the diffusion constant, *τ* is the time lag, and the term 2*σ*^*2*^ represents the mean localization error in *x* and *y* ^26^. The blue curve in **Fig. 2b** presents the MSD, which displays a systematic and non-trivial deviation from the expected behavior of normal diffusion illustrated by the black straight line. The observed *MSD* shows a substantial tendency toward sub-diffusion and, hence, the existence of binding or hindering interactions ^27^. We point out that the localization precision in most published works is only determined categorically for the microscope (see e.g. Refs. [^3,28^]), but in our experiments it was assessed for each frame. We also emphasize that inclusion of the positional uncertainty *σ* is important for a correct assessment of the MSD ^26^, and especially in high-speed imaging where the mean square displacement becomes comparable to the localization precision.

A topic of great interest regarding diffusion in the plasma membrane has been the occurrence of transient nanoscale phenomena such as lipid rafts or protein clustering ^25,29^. Efforts to identify local confinements typically fall within two approaches, both based on assumptions. One strategy builds on *a priori* knowledge of the effective diffusion constant and infers confinement when the displacement falls short of the expected travel ^30,31^. An alternate approach considers a specific geometry and fits the measured *MSD* to a model derived for that scenario, e.g., hopping across semi-permeable barriers of square confinements ^32^ (see examples in Ref. [^33^]). We can afford not to make any assumptions about the nature of the diffusion or its geographic landscape because the very large number of iSCAT frames and the localization precision in each frame make it possible to segment each trajectory into small windows while ensuring robust statistics.

We examine local deviations from normal diffusion by considering

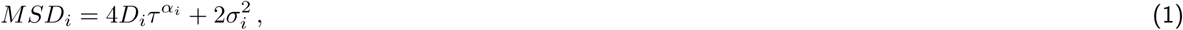

for the *i*-th rolling temporal window. This formulation provides a convenient measure for the nature of diffusion: while *α* is unity for normal diffusion, *α <* 1 indicates subdiffusion, and *α >* 1 is a sign of superdiffusion ^34^. In **Fig. 2c**, we map *α*_*i*_ in a color-coded fashion, indicating varying degrees of middling and strong confinement. An example of the latter is a region marked with the dashed circle which will be explored further below. To facilitate a quantitative insight into the overall trajectory, we also plot *α*_*i*_ as a function of time in **Fig. 2d**, whereby the gray regions depict the uncertainty. Interestingly, the regions where *α*_*i*_ clearly deviates from unity seem to be locally confined to a few tens of nanometers and within time intervals of tens to hundreds of milliseconds. An example corresponds to the region marked with * in (**c**) where *α*_*i*_ *<* 0.75. We note that our findings do not depend on the length of the rolling windows.

Another approach for gauging the occurrence of diffusive barriers and confinements is to consider the degree of directional correlation (*C*_*i*_) between two vectorial steps across a time lag ^35^. A truly random process would possess no directional correlative memory (median *C*_*i*_ = 0), whereas negative or positive values would indicate reflections or forwardly push, respectively. **Figure 2e** shows the 2D map of *C*_*i*_, and in **Fig. 2f** we plot the correlation coefficients over sliding windows of 500 frames with the same temporal lag as used for determining *α*_*i*_. We find local variations on a baseline of *C* = − 0.15 while comparison of **Fig. 2e** with **Fig. 2c** and **Fig. 2f** with **Fig. 2d** confirms a correspondence between the areas of pronounced sub-diffusion (*α*_*i*_ *<* 1) and strong backlash (*C*_*i*_ *<* 1).

As a third way of discovering heterogeneities, we assessed the popularity of each trajectory pixel in space. Here, we introduce the concept of accumulated temporal occupancy map (ATOM) wherein we divide the lateral plane of the trajectory into nanometer-sized bins as small as the localization precision and count the occurrence of the particle in each bin. The areas of extended residency in **Fig. 2g** identify nanoscopic persistent structures arranged in loops and whirls (see **i**-**iii**) as well as a patch extending over approximately 150 nm marked by *, whereby the latter corresponds well to the marking in **Fig. 2c**. We observed a minimal lifetime of 250 ms (5,000 frames) for this feature, which might portray a pre-endocytic step to membrane invagination (we will return to this discussion). Overall, the observed inhomogeneities strongly suggest that protein diffusion is affected by cellular sub-structure.

### Diffusion along a filopodium over minutes

An important advantage of iSCAT microscopy is its ability to record very long events. We now apply this feature together with the 3D imaging capability of iSCAT to follow EGFRs on a filopodium. Filopodia are rod-like cellular protrusions containing bundles of actin filaments with diameter of 100-300 nm and up to 10 *µ*m in length. They act as sensors for mechanical stimuli, putative attachment sites, chemoattractants or repellents in the surroundings of the cell. Filopodia also provide cellular signaling sites enriched in growth factor receptors ^36,37^, and indeed, it has been shown that ligand binding and EGFR activation on filopodia occur preferentially at low EGF concentrations. Activation is then followed by association with the actin filaments and retrograde transport of EGFR to the cell body, where the receptors are endocytosed ^38,39^. We now show how iSCAT microscopy gives new insight into the nanoscopic details of this process. To set the stage, in **Fig. 3a** we display a transmission electron microscope snapshot of a typical filopodium from a HeLa cell, including an EGF-GNP.

**Figure 3:**
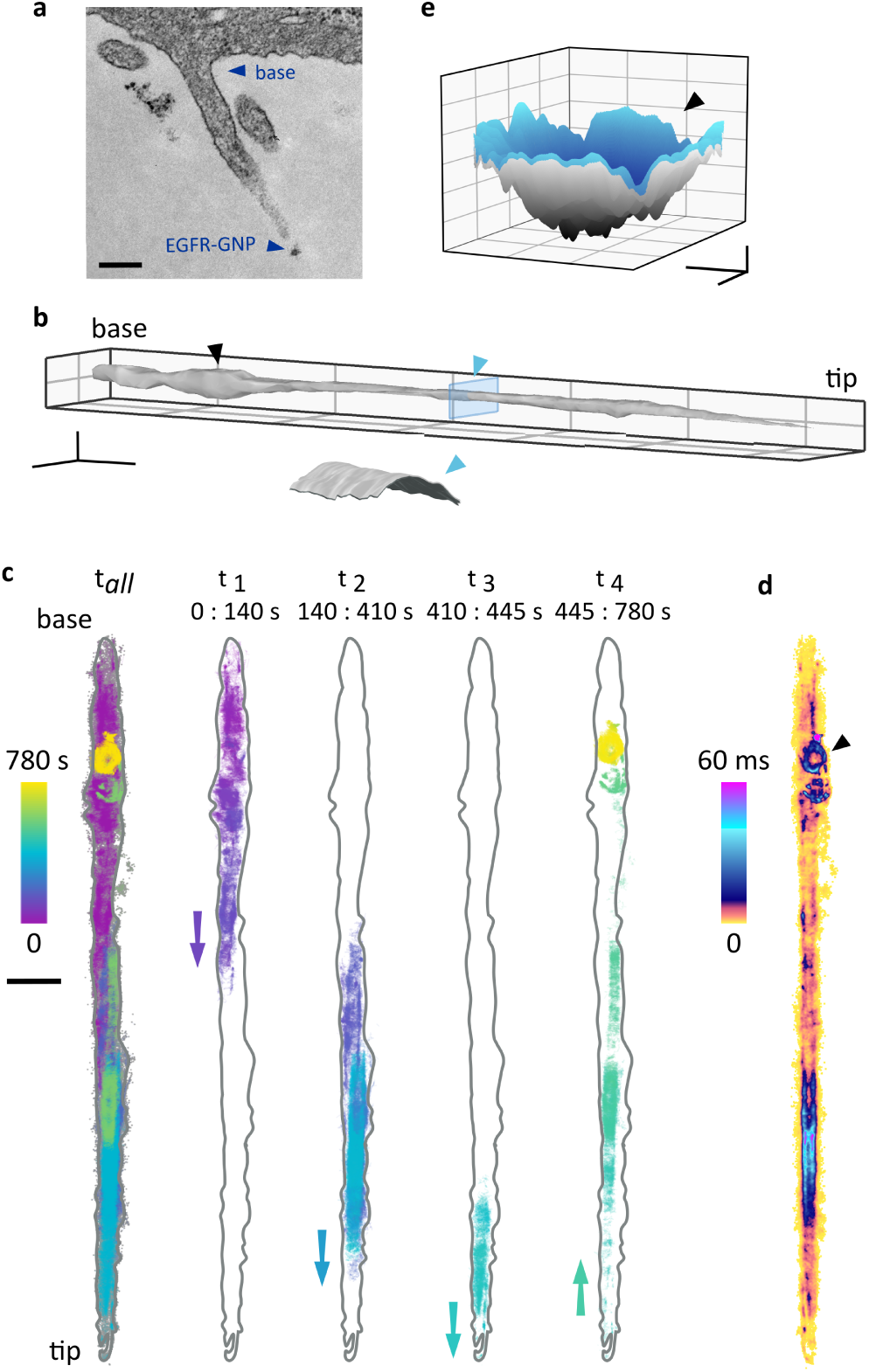
Long journey of an EGFR: Mapping a filopodium by a nano-rover. (a) Electron microscopy image of a HeLa filopodium including an EGF-GNP probe near the tip. Scale bar indicates 200 nm. (b) Filopodium surface reconstructed from the 780,000 recorded trajectory positions. Scale bars denote 1 *µ*m (*x*,*y*) and 200 nm (*z*). Here, we reduced the imaging speed to 1,000 fps (localization error *σ*_*x,y*_ = 2 nm) to avoid complications with capture and storage of large data volumes. The inset shows the cross section of the blue slice and points to a cylindrical surface of diameter 150 nm after accounting for the size of the GNP. (c) The 13 minute trajectory (left) and a breakdown of the trajectory into four subsequent pieces, revealing walk to the tip and return to the base. Scale bar 1000 nm. (d) ATOM plot of the trajectory. We note that this figure was corrected for filopodium drift over 13 min while the data in (c) showed the raw trajectory before correction. (e) A surface interpolation of the trajectory from the final 80 s. The ring-like confinement in the final phase of the trajectory is found to be a 3D pit (marked with black triangle). Scale bars denote 100 nm (*x,y*) and 50 nm (*z*).

In **Fig. 3**, we present data obtained from an iSCAT video recorded over the course of 13 minutes. First, we analyse the 3D trajectory of the EGFR-GNP, whereby the very large number of trajectory points makes it possible to interpolate among them and create the topography of the filopodium surface as displayed in **Fig. 3b**. The 3D locations of the recorded positions map to a cylinder-like shape with a diameter, which when corrected for the diameter of the GNP amounts to about 150 nm. The cut through the filopodium displayed in the inset clearly shows that the probe has explored the outer surface. This provides an example, where a GNP serves as a nano-rover for mapping the topology of cellular structures, while its 3D coordinates are registered via iSCAT.

In **Fig. 3c**, we examine the full trajectory of the EFGR-GNP in more detail by visualizing sections from different time intervals. The color-coded temporal evolution of the trajectory reveals a directionality accompanied by a longitudinal dithering behavior. We find that the receptor initially travels toward the tip of the filopodium over a period of 445 s, followed by retrograde trafficking back towards the cell body where it is trapped in a confinement. This observation supports the literature reports of directed transport of receptor-bound EGF along filopodia ^38^.

To learn more about the phenomena at work, in **Fig. 3d** we plot the trajectory ATOM. We find that the occupancy of the filopodium tip region is short compared to the residence time along the shaft. ATOM data also reveal extended residency in a circular patch and a ring-like structure towards the end of the trajectory (marked with a black triangle). Interestingly, the 3D representation of this confinement in **Fig. 3e** indicates a pit-like topology, which would be consistent with a pre-endocytic step of membrane invagination.

### A closer look at examples of confined diffusion and evidence for uptake

Another asset of iSCAT stems from the linearity of the scattering signal and lack of saturation, which limits the speed of fluorescence imaging. High-speed microscopy is not only necessary for obtaining high-resolution temporal information, but it also circumvents blurring effects in the spatial localization of nanoparticles ^40^. **Figure 4a** portrays a bounded trajectory within a disk of 500 nm diameter over 2.6 s, an observation that is consistent with long-time confinements reported for EGFR^3,41^.

**Figure 4:**
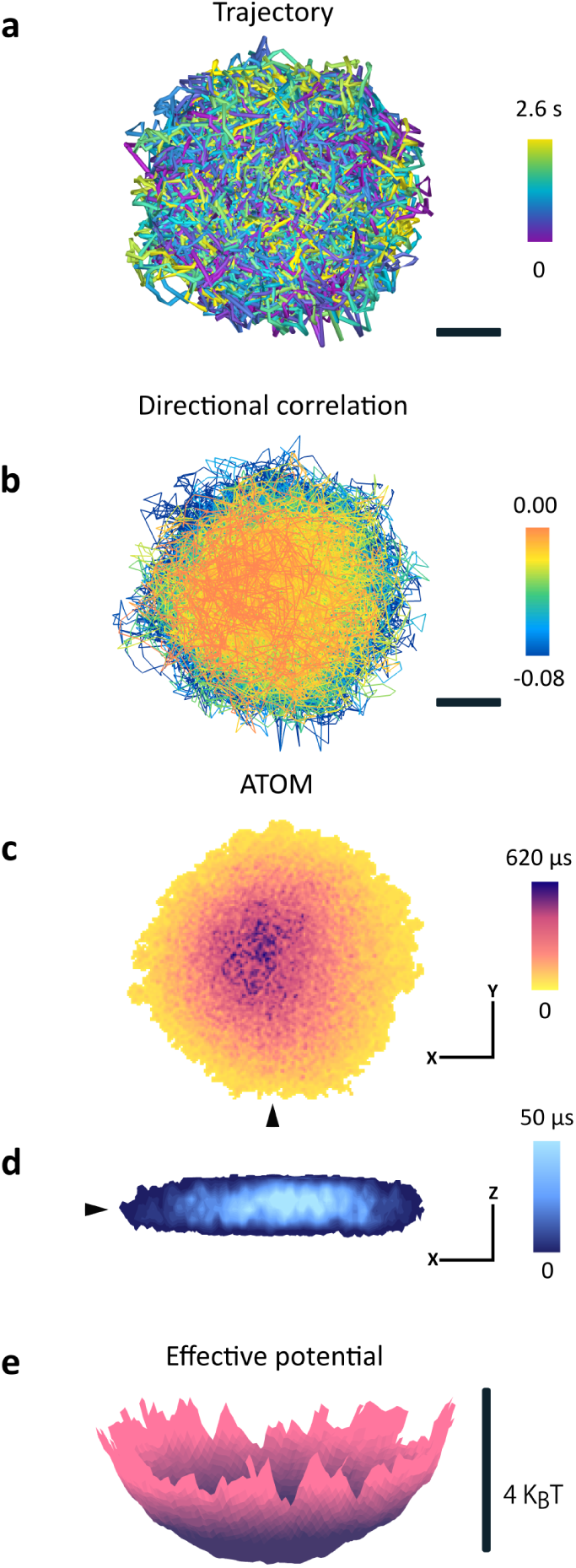
Confined diffusion recorded at 30,000 fps. (a) Lateral trajectory color coded for time. Scale bar 100 nm. A lower temporal sampling of this confinement would have under-estimated the extent of bounding ^42^. (b) Directional correlation of the trajectory (using a time lag of five frames), showing partially hindered diffusion with a propensity for freer diffusion in the centre. (c) ATOM plot of (a). (d) A cut through the 3D-ATOM plot along the line of the black triangle in (c), shows that occupancy favors an inner most disk-like region. Axes denote 100 nm in both (c) and (d). Color bars encode the residency times. (e) Conversion of the temporal 2D occupation from (c) into an effective potential distribution.

The directional correlation *C*_*i*_ and the 3D-ATOM in **Fig. 4b-d** provide further evidence for confinement in all dimensions. In particular, the lower values of *C*_*i*_ on the outer part of the trajectory clearly visualise an act of confinement, which is supported by a propensity for the protein to be found in the center of the ATOM plot. Furthermore, we used the ATOM data to compute the normalized probability amplitude of spatial occupation *P* (*x, y*) and deduced an effective bounding potential according to *U* = *-k*_*B*_*T* log(*P* (*x, y*)) ^42^. As displayed in **Fig. 4e**, we find a concave functional form with depth of *≈* 4 *K*_*B*_*T*, consistent with the results of Ref. [^42^].

In **Fig. 5a**, we present another 3D trajectory recorded at a very fast rate of 66,000 fps with a short exposure time of 10 *µ*s and duration of 3.5 s, while maintaining a high spatial precision of *σ* = 4 nm. To portray the axial variations of the particle coordinates in a quantitative fashion, in **Fig. 5b** we plot the height as a function of time with the same color code as in (**a**). We see that the protein undergoes a substantial out-of-plane motion of up to 200 nm over 2 s. For comparison, we also plot the height variation of the membrane as a whole during the same time, deduced by analyzing the iSCAT background.

**Figure 5:**
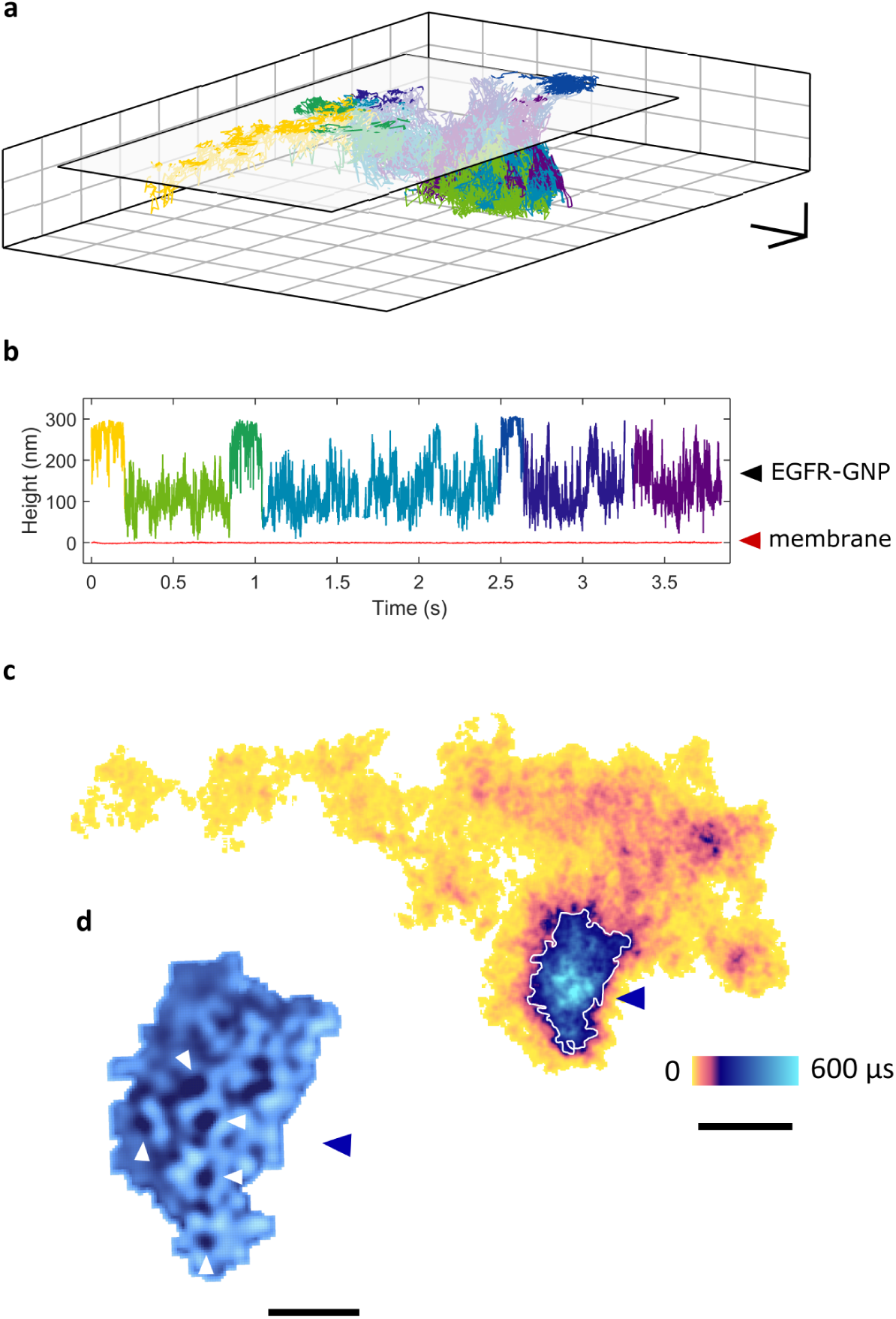
Ultra-highspeed 3D tracking at 66,000 fps. (a) 3D trajectory over 3.75 s with *σ*_*x,y*_ = 4 nm. The initial surface of diffusion, marked with a white plane, sits above a depressed valley region. Scale bar denotes 200 nm (*x*,*y*) and 100 nm (*z*). Color code represents time according to the representation in (b). (b) Height of probe as a function of time emphasizes substantial axial motion of the EGFR-GNP probe. The instantaneous background fluctuations of cell membrane (red) corresponding to a time-averaged value of 0.5 nm show clearly uncorrelated motion to that of the EGFR-GNP probe. (c) The ATOM plot. A patch of extended occupation (blue) within the topological valley region is easily identified. Scale bar 250 nm. (d) Upon closer inspection after removal of a background, a network of 25 nm rings (white pointers) becomes evident. Scale bar denotes 50 nm.

Next, in **Fig. 5c** we display the ATOM plot of this trajectory, which again, indicates a good deal of heterogeneity in residence times. A notable feature now is the depressed valley region, where the protein spends more time. **Figure 5d** shows a zoom into this patch, revealing circular features of 25 nm diameter arranged in a quasi-lattice. The overall size of the confinement and the dimensions of the underlying network of rings surveyed by the nano-rover are consistent with the features of a clathrin-coated lattice or pit ^43^. It is known that the assembly of a clathrin coat on the inner surface of the plasma membrane results in the formation of clathrin-coated patches and can occur spontaneously or in a cargo-triggered manner. Once associated with a cargo protein, basket-like structures grow to form clathrin-coated pits (CCP), inducing membrane invagination. CCPs are reported to cover about 1.5% of a HeLa cell surface, accompanied by flat clathrin lattices that can cover up to 8% of its inner membrane ^44^.

EGFR is expected to undergo clathrin-mediated endocytosis (CME) in HeLa cells when stimulated by a low concentration of EGF (order of pM) ^45^, a phenomenon that we confirmed through fluorescence confocal microscopy. In **Figures 3**, **4** and **5**, we showed that fast iSCAT microscopy can reveal intricate 3D features that hint toward endocytotic events. We now provide one more such evidence.

**Figure 6a** displays a conventional 2D map of a confined trajectory over a duration of 1 s, while its 3D smoothed map in **Fig. 6b** illustrates a bowl-like topology with diameter of 150-200 nm (after correction for the size of the GNP) and lasting for tens of seconds, consistent with the literature knowledge on CME^46–48^. The fact that the GNP maps a curved surface suggests that the EGFR is not only engulfed by the pit but it also stays attached to and explores its inner surfaces. To verify that our EGFR-GNP entities are captured in CCPs, we also performed electron microscopy on our samples. **Figure 6c** presents an example, where EGFR-GNPs (pink triangle) are located in a CCP (white triangle) and a vesicle (yellow triangle).

**Figure 6:**
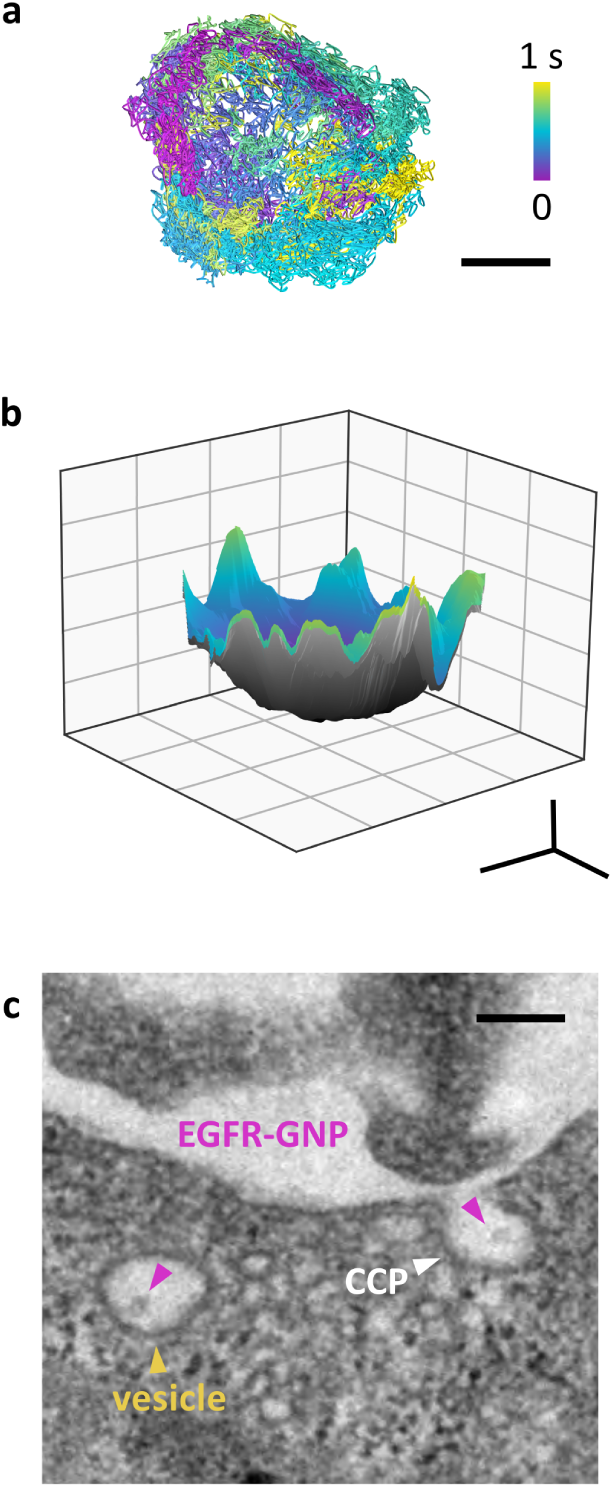
Confined diffusion within a pit. Tracking a protein at high speed of 20,000 fps (17.5 *µ*s exposure time). (a) Lateral trajectory of the diffusive path. The fast motion of EGFR is confined to a circular pit of 200 nm diameter. Scale bar 100 nm. (b) The 3D projection of (a) maps to a bowl-like surface. Scale bar denotes 100 nm (*x,y*) and 20 nm (*z*). (c) Ultra-thin (50 nm) electron microscopy section image of a HeLa cell confining an EGFR-GNP probe (pink triangles) to a clathrin-coated pit (white triangle) and a vesicle (yellow triangle). Scale bar 200 nm. Cell fixation occurs near-immediately after addition of EGF-GNP probes, consistent with the time frame of (a,b).

We remark that the finite GNP diameter of 40-50 nm seems not to be prohibitive in our studies. In fact, this is not surprising given the much larger size of the CCP. Furthermore, considering that EGFR is expected to be engulfed in small lipid vesicles of size comparable to our GNPs after its uptake ^47,48^, the GNP size does not necessarily impose extra space constraints. Furthermore, the size of putative CCPs and internalized vesicles observed in iSCAT match those seen in our TEM images with and without EGFR-GNPs present, indicating lack of systematic artefacts. We also note that EGFR is expected to undergo multimerization in the endocytotic process^49^. Although our current measurements are not able to examine this matter, our findings have been consistent with the existing knowledge about the interaction of EGFR with the plasma membrane and clathrin.

## Discussion and outlook

After several decades of hard work and clever experimentation with a wide range of biochemical, analytical and imaging techniques, our knowledge of the cellular plasma membrane continues to be limited. In particular, it is still unclear how lipids, proteins and small molecules within the membrane interact with each other and with their neighbors such as other proteins, the cytoskeleton or extracellular matrix. In the case of transmembrane proteins, different models and hypotheses have been put forth to explain the observed inhomogeneities in their diffusion behavior. In our current work, we have not attempted to verify or refute any of the existing hypotheses, which are possibly still incomplete due to lack of differentiated data. Rather, we present intriguing examples where the combination of very high spatial and temporal resolution in iSCAT microscopy provides unprecedented details about sub-diffusion, nanoscopic confinement and 3D contours of filopodia and clathrin structures. These findings help establish iSCAT as a powerful tool for quantitative studies in cell biology.

In future, we plan to combine iSCAT with *in-situ* super-resolution fluorescence microscopy to correlate iSCAT trajectories of proteins and viruses with the structures of other entities such as actin filaments, microtubuli, clathrin or dynamin. For example, it would be interesting to investigate hypotheses such as hopping or corralling by the actin mesh ^25^ as well as temporal and spatial details of endocytosis ^43^. A particularly promising application of our methodology will be in understanding the life cycle of viruses, which provide a large iSCAT signal without the need for an external label ^9,28,50^.

## Materials & Methods

### Synthesis of EGF-GNP probes

Gold nanoparticles functionalized with monoclonal anti-biotin (British Biocell International) were conjugated at a molar ratio of 1:1 with biotinylated-EGF (ThermoFisher) to a concentration of 0.66 nM. A 1% (w/v) biotin-free PBS-BSA solution was used as a buffer. The solution containing EGF-GNP was agitated for several hours at 37 °C to assist conjugation, purified through a NAP-5 column (GE Healthcare) and diluted to a final concentration of 0.1 nM.

### Cell culture

HeLa cells were grown in Dulbecco’s Modified Eagle Medium (Gibco, Invitrogen) supplemented with 10% fetal calf serum (Life Technologies) in a humidified atmosphere at 37 °C and 5% CO2. For measurement, the cells were plated onto a glass-bottomed sample dish (Ibidi GmBH) and grown to 70% confluency. Prior to measurement, each dish was rinsed twice with warmed PBS-BSA solution and serum starved for several hours and finally rinsed again with warmed PBS-BSA. When placed on the microscope, a buffer of Leibovitz’s L-15 Medium (1.5 ml Gibco, Invitrogen) was added and the culture was held at 37 °C via a home-built objective heater. A pipette was used to deliver 10 *µ*l of the EGF-GNP to a local region of the culture with observation beginning immediately thereafter.

### Immunofluorescence

HeLa cells were seeded on fibronectin-coated coverslips inserted into 24-well plates (Sarstedt) to a confluency of 1×10^5^ cells per well. 24 h after seeding, the cells were serum starved (DMEM, 0.1% FCS) for 16 h. The plates were chilled on ice for 1 h and then stimulated for 4 min at 37 °C with either the control medium, 16 nM EGF or 1 pM EGF-GNP with additional pre-warmed DMEM (with 0.1% FCS). After stimulation the samples were washed twice with ice-cold PBS (Gibco, Life Technologies) and fixed with 4% paraformaldehyde for 15 minutes. Cells were then permeabilized with 0.1% TritonX-100 (Carl Roth) and blocked with 10% horse serum/PBT (Phosphate-Buffered Saline supplemented with 0.1% Tween 20). EGFR was detected using a specific primary antibody rat anti-EGFR (abcam ICR10, 1:1000) and a secondary anti-rat-Cy3 antibody (Jackson Immuno Research, 1:400). Clathrin or caveolin was visualized using specific antibodies: rabbit-anti-clathrin (proteintech, 1:200) or rabbit-anti-caveolin (proteintech, 1:500) and a secondary anti-rabbit-Alexa488 antibody (ThermoFisher, 1:1000). Nuclei were stained with DAPI. The samples were mounted in Mowiol and imaged with a confocal laser scanning microscope.

### Preparation of cell-lysates and western blotting

HeLa cells were seeded at a confluency of 3×10^5^ into a 6-well plate (Sarstedt), cultured for 24 h followed by incubation over-night in starvation medium (DMEM, 0.1% FCS). Stimulation was performed using 10 pM or 100 pM EGF, EGF-GNP or control GNP respectively for 2 min. Cells were lysed in 250 *µ*l of 20 mM HEPES (pH 7.4), 150 mM NaCl, 0.5% NP-40 supplemented with protease inhibitor and phosphatase inhibitor cocktails (Roche) at 4 °C. Lysates were cleared by centrifugation for 10 min at 4 °C and 16,000 g. The total amount of protein was determined using a BCA assay (Applichem). Equal amounts of protein per sample were mixed with SDS sample buffer and denatured for 5 min at 95 °C. The proteins were separated by gel-electrophoresis (10% polyacrylamide gel) and transferred onto a PVDF membrane (Roche) using a semi-dry blotting system (BioRad). Primary antibodies used were rabbit-anti-EGFR (Cell Signalling Technologies (D38B1), 1:1000), mouse-anti-GAPDH (proteintech, 1:2000), rabbit-anti-Erk1, 2 (Cell Signaling Technologies, 1:1000), rabbit-anti-pErk (Promega, 1:2000). Secondary antibodies anti-mouse-Ap and anti-rabbit-Ap (Cell signalling Technologies, 1:3000).

### Electron Microscopy

HeLa cells were seeded at a confluency of 5×10^5^ into 4-well-lumox chamber slides (x-well-Lumox, Sarstedt). Stimulation was performed as described for immunofluorescence using 1 pM EGF-GNP. After stimulation the cells were rinsed twice with ice-cold PBS and fixed with 25% glutaraldehyde and 4% paraformaldehyde in 0.1 M cacodylate buffer pH 7.4 for 15 min at 37 °C. Cells were washed once with PBS and fixed again with 4% osmium tetroxide in 0.1 M cacodylate buffer pH 7.4 for one hour in the dark. Samples were dehydrated by an ascending acetone series and embedded in an Epon-Acetone-100% mix (1:1) over night at 60 °C. Samples were embedded in pure Epon on the next day. Ultrathin sections of 50 nm were cut at an ultra-microtome (Ultra cut E, Reichert & Jung). Thin sections were mounted on a copper grid and counter stained with a uranyl acetate / lead-cytrate-mix. Samples were imaged on a Transmission Electron Microscope (Zeiss).

### PSF extraction and 3D localization

A typical iSCAT image in our work contains contributions from scattering by the particle and the cell. To extract the PSFs of the probe in different frames of a recorded video, we first fit a 2D Gaussian function to the image of the GNP in one of the video frames which possesses a sufficiently large positive contrast to enable a precise determination of the centroid position. Next, we compute the radial median intensity about this centroid and use this information to reconstruct the PSF for that frame (see **Fig. 1d**). This approach is made possible by the strong circular symmetry of the PSF against the asymmetric fluctuations of the speckle background.

The PSF (*PSF*_*t*1_) extracted from video frame *V*_*t*1_ is cross-correlated with the next time-subsequent frame *V*_*t*2_ to determine the probe location in the latter. Here, we compute a 2D intensity cross-correlation following *O*(*u, v*) = Σ_*x, y*_ *PSF*_*t*1_(*x, y*)*V*_*t*2_(*x − u, y − v*). Next, a Gaussian function is fitted to the maxima in the resultant cross-correlation product *O*, and the uncertainty in the lateral position of the localized probe is computed from this fit. This approach is repeated throughout the entire video and is robust against changes in PSF contrast polarity. We point out that this algorithm relies on our fast imaging frame rate, which avoids large jumps in the trajectory. The concentric ring pattern of the PSF enables determination of the axial position of the probe.

## Acknowledgements

This project was funded by an Alexander von Humboldt professorship, Max Planck Society and the Research and Training Grant 1962, ‘Dynamic Interactions at Biological Membranes’ of the German Research Foundation (DFG). RWT acknowledges an Alexander von Humboldt fellowship. VR and AS were also supported by a grant from DFG (Grant SCHA965/9-1). We thank Simone Ihloff for support in cell culturing, Claudia Obermeier for preparing ultra thin sections (TEM), Benjamin Schmid (Optical Imaging Center Erlangen, OICE) for support in co-localization analyses and Vasily Zaburdaev for insightful discussions regarding statistical analysis of diffusion.

## Competing Interests

The authors declare no competing interests.

